# Phylogenomics reveals that *Asaia* symbionts from insects underwent convergent genome reduction, preserving an insecticide-degrading gene

**DOI:** 10.1101/2020.10.30.363036

**Authors:** Francesco Comandatore, Claudia Damiani, Alessia Cappelli, Paulo Ribolla, Giuliano Gasperi, Francesco Gradoni, Gioia Capelli, Aurora Piazza, Fabrizio Montarsi, Maria Vittoria Mancini, Paolo Rossi, Irene Ricci, Claudio Bandi, Guido Favia

**Author notes:** Francesco Comandatore and Claudia Damiani contributed equally to this work, *Author order was determined both alphabetically and in order of increasing seniority*. Corresponding author: Guido Favia. School of Biosciences & Veterinary Medicine, University of Camerino, CIRM Italian Malaria Network, Camerino (MC), Italy.

## Abstract

The mosquito microbiota is composed by several lineages of microorganisms whose ecological roles and evolutionary histories have yet to be investigated in depth. Among these microorganisms, *Asaia* bacteria play a prominent role, given its abundance in the gut, reproductive organs and salivary glands of different mosquito species, while its presence has also been reported in several other insects. Notably, *Asaia* has great potential as a tool for the control of mosquito-borne diseases. Here, we present a wide phylogenomic analysis of *Asaia* strains isolated from different species of mosquito vectors and from different populations of the Mediterranean fruit fly (medfly) *Ceratitis capitata*, an insect pest of worldwide economic importance. We show that phylogenetically distant lineages of *Asaia* experienced independent genome reductions, despite following a common pattern, characterized by the early loss of genes involved in genome stability. This result highlights the role of specific metabolic pathways in the symbiotic relationship between *Asaia* and the insect host. Finally, we discovered that all but one of the *Asaia* strains included in the study possess the pyrethroid hydrolase gene. Phylogenetic analysis revealed that this gene is ancestral in *Asaia*, strongly suggesting that it played a role in the establishment of the symbiotic association between these bacteria and the mosquito hosts. We propose that this gene from the symbiont contributed to initial pyrethroid resistance in insects harboring *Asaia*, also considering the widespread production of pyrethrins by several plants.

**Importance:** We have studied genome reduction within several strains of the insect symbiont *Asaia*, isolated from different species/strains of mosquito and medfly. Phylogenetically distant strains of *Asaia*, despite following a common pattern involving the loss of genes related to genome stability, have undergone independent genome reductions, highlighting the peculiar role of specific metabolic pathways in the symbiotic relationship between *Asaia* and its host. We also show that the pyrethroid hydrolase gene is present in all the *Asaia* strains isolated except for the South American malaria vector *An. darlingi*, for which resistance to pyrethroids has never been reported, suggesting a possible involvement of *Asaia* in determining the resistance to insecticides.

## Introduction

Symbiosis shaped the evolution animals and plants, providing new adaptative traits to multicellular organisms and increasing their capability to create and colonize novel ecological niches (1). On the other hand, microbial symbionts have also been subjected to drastic functional and metabolic modifications, associated with a reduction of the genome size (2). The phenomenon of bacterial genome reduction has been shown to correlate with genome-wide loss of genes. This is particularly well described in obligate bacterial endosymbionts of insects, such as *Blattabacterium cuenoti*, *Buchnera aphidicola* and *Carsonella ruddii* (3,4).

We recently provided the first evidence of genome reduction in *Asaia* (5), a major component of the gut microbiota in several insects, including relevant human disease vectors (6). Indeed, *Asaia* has been detected in *Anopheles, Aedes* and *Culex* mosquito species, which are known to transmit several pathogens to humans, including the etiological agents of malaria, dengue, yellow fever, zika, chikungunya and some filarial diseases (6–10). Notably, *Asaia*, apart from its well-established role as a main component of the insect microbiota, can also be regarded as an environmental bacterium, generally associated with sugar-containing water pools, such as those in flowers and leaf axils in several plants (11).

In our previous studies we showed that *Asaia* influences both the development and the longevity of *Anopheles stephensi* mosquitoes (12,13). Moreover, experimental evidence indicates that *Asaia* affects the immune-related gene expression in mosquitoes (13–15). Overall, the collective evidence suggests that *Asaia* plays a crucial role in mosquito development, physiology and survival.

Interestingly, *Asaia* has also been detected in insect pests and vectors of agricultural relevance, such as *Scaphoideus titanus* and the white-backed planthopper, *Sogatella furcifera* (6,16). In this context, we have recently detected *Asaia* in different strains of *Ceratitis capitata* (Damiani et al, in preparation), a major pest threat to the agriculture in several geographical regions world-wide (17).

Our previous study showed that *Asaia* from *Anopheles darlingi*, a South American malaria vector (5) underwent a significant reduction both in genome size and in gene content. Consequently, *Asaia* was proposed as a novel model to investigate genome reduction, approaching this phenomenon within a single bacterial taxon that includes both environmental and insect-associated strains or species.

Here we present a comprehensive comparative genomics analysis of different *Asaia* strains isolated from different sources including several mosquito species, different populations of *C. capitata*, and environmental samples. Our results revealed substantial changes in genomic content across different *Asaia* lineages, showing independent genome reduction processes that occurred through a common pattern. Furthermore, we found that all but one of the *Asaia* strains examined harbour the pyrethroid hydrolase gene, which is likely to confer resistance to pyrethroids, insecticides produced by plants and commonly used in pest treatments (18).

## Results and Discussion

A total of 17 novel genomes were obtained from *Asaia* strains, from mosquitoes and *Ceratitis capitata;* these were compared with the 13 *Asaia* genomes already available in the data base, obtained from stains isolated from mosquitoes and from environmental samples (see Table S1 in the supplemental material).

Orthologous analysis allowed the identification of 612 core genes for an amino acidic concatenate, sized 183,373 base-pairs. The obtained Maximum Likelihood phylogenetic tree is shown in Fig. 1. Average Nucleotide Identity (ANI) analysis clustered *Asaia* strains into six sub-groups, corresponding to the previously described *Asaia* species (Fig. 1). An integrative analysis has been performed with a larger number of outgroups, confirming the divergent clustering of *Asaia* while retaining the six sub-groups (see Fig. S1 in the supplemental material).

**Fig. 1.**
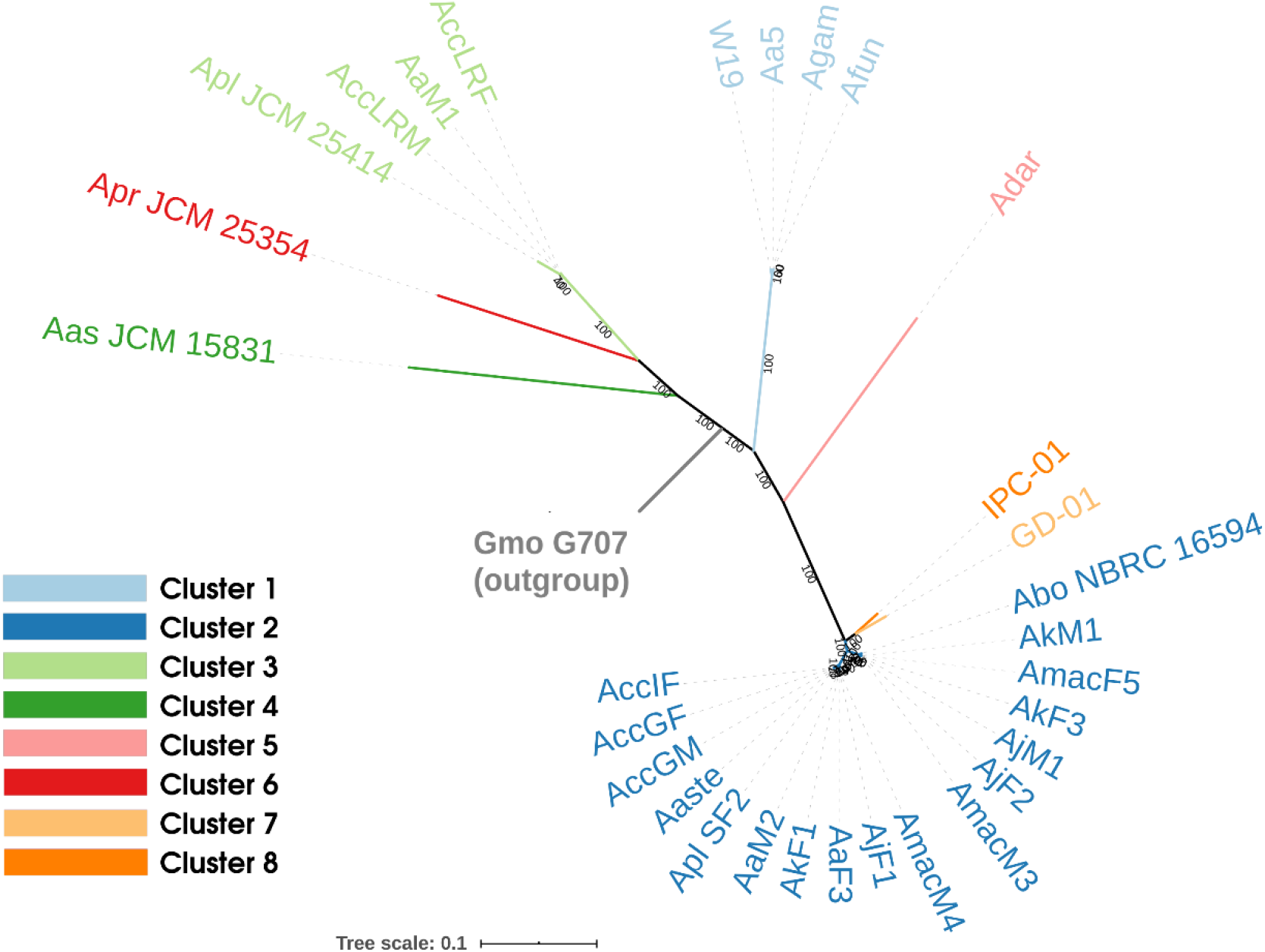
Maximum Likelihood phylogenetic tree. Maximum Likelihood phylogenetic tree obtained using the 612 core genes among the 30 *Asaia* spp. strains included in the study and the *Gluconobacter morbifer* strain G707 as outgroup. Tree leaves and the relative monophyletic clades colors represent the clusters identified on the basis of Average Nucleotide Identity (ANI) with a cut-off of 85%. Bootstrap support values are reported on the branches of the tree.

It is worth nothing that the current taxonomic classification of the *Asaia* strains is currently debated, indeed most of the sequenced strains are not officially assigned to specific *Asaia* species. The ANI-based analysis with 95% threshold is often used to describe bacterial species (19), suggesting that the “clusters” we found may correspond to *Asaia* species coherently we retain the term “cluster”.

The variance underlying the specific clustering of the different *Asaia* strains reflects the existence of a differential gene composition, thus suggesting an evolution that occurred through independent genomic reduction/variation pathways. The principal coordinates analysis (PCoA) on gene presence/absence across the different *Asaia* strains analysed, clearly indicate that these differences are not solely attributable to ecological adaptations (Fig. 2) but they should be interpreted as a consequence of the speciation process.

**Fig. 2.**
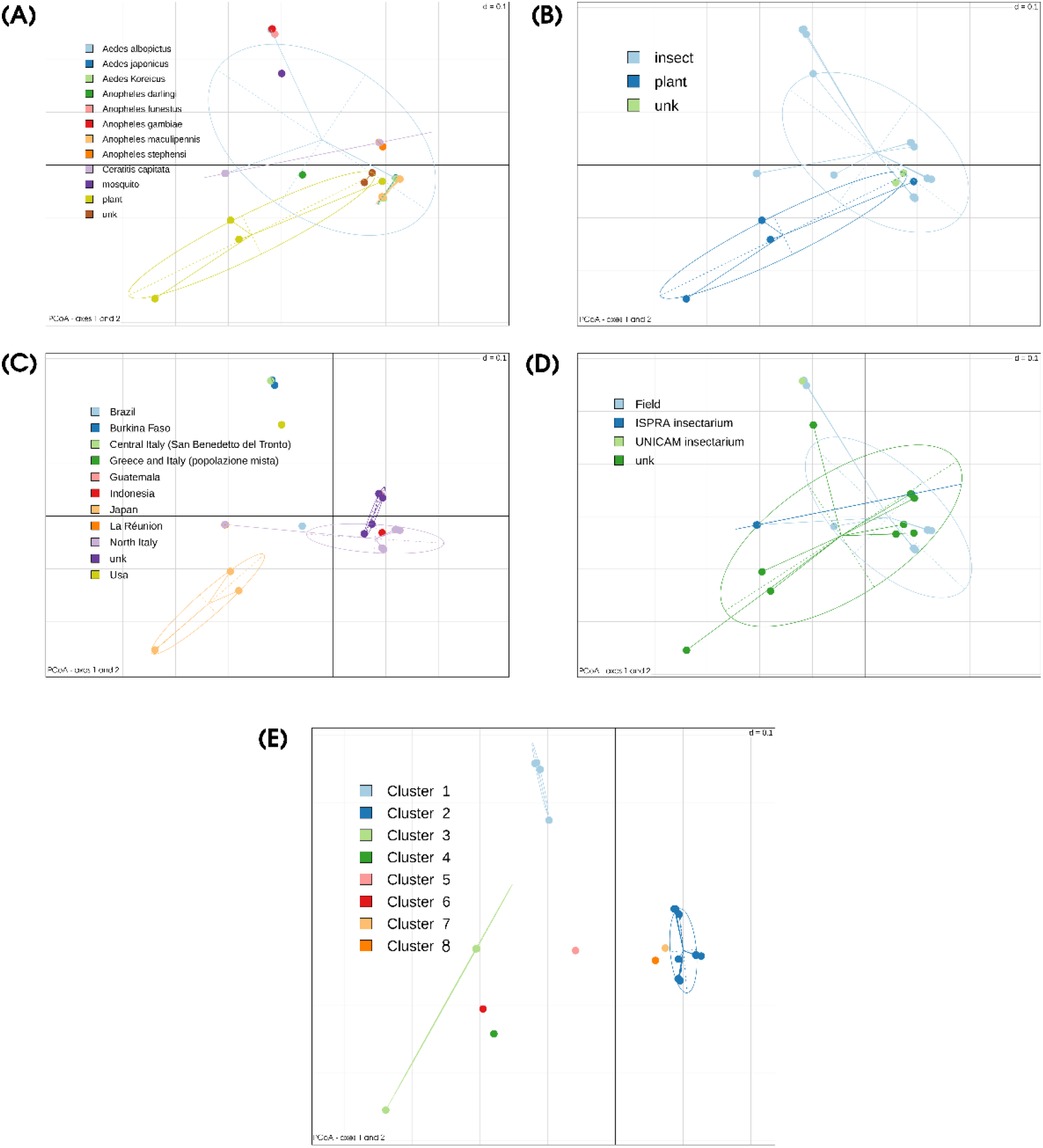
Principal Coordinates analysis. Bi-Dimensional Principal Coordinates Analysis plots generated on the basis of gene presence / absence (explained variance 24,54% on the x-axis and 16,10% on the y-axis). Each dot of the plots represents one of the 30 *Asaia* spp. strains included in the study. Plot (a): each color corresponds to a single host species; plot (b): *Asaia*-hosts are grouped in the categories insects, plants and unknown; plot (c) *Asaia*-hosts are grouped on the basis of the sampling geographic location; plot (d) *Asaia*-hosts are grouped on the basis of the sample origin; plot (e) *Asaia*-hosts are grouped on the basis of clusters identified using Average Nucleotide Identity (ANI) with a cut-off of 95%.

Even if the presented evidence suggests for an independent evolution of the different *Asaia* strains, we found that genomic erosion tends to preserve some pathways, while others erode more easily in all strains (Fig. 3). In particular, only the “Replication, recombination and repair” (L), “Transcription” (K) and “Cell Motility” (N) Cluster of Orthologous Groups (COGs) pathways showed a clear erosion trend (Fig. 3a), while several others resulted preserved (Fig. 3b and see Table S2 in the supplemental material). Notably, most of the preserved pathways are involved in essential metabolic functions of the bacteria (e.g. transport and metabolism of carbohydrate, amino acid, nucleotide, co-enzyme, as well as membrane biogenesis and energy production). This clearly suggests that the erosion process tends to preserve genomic stability *tout-court* pathways rather than being subjected by functional/ecological adaptation processes, at least in the early stages of reduction.

**Fig. 3.**
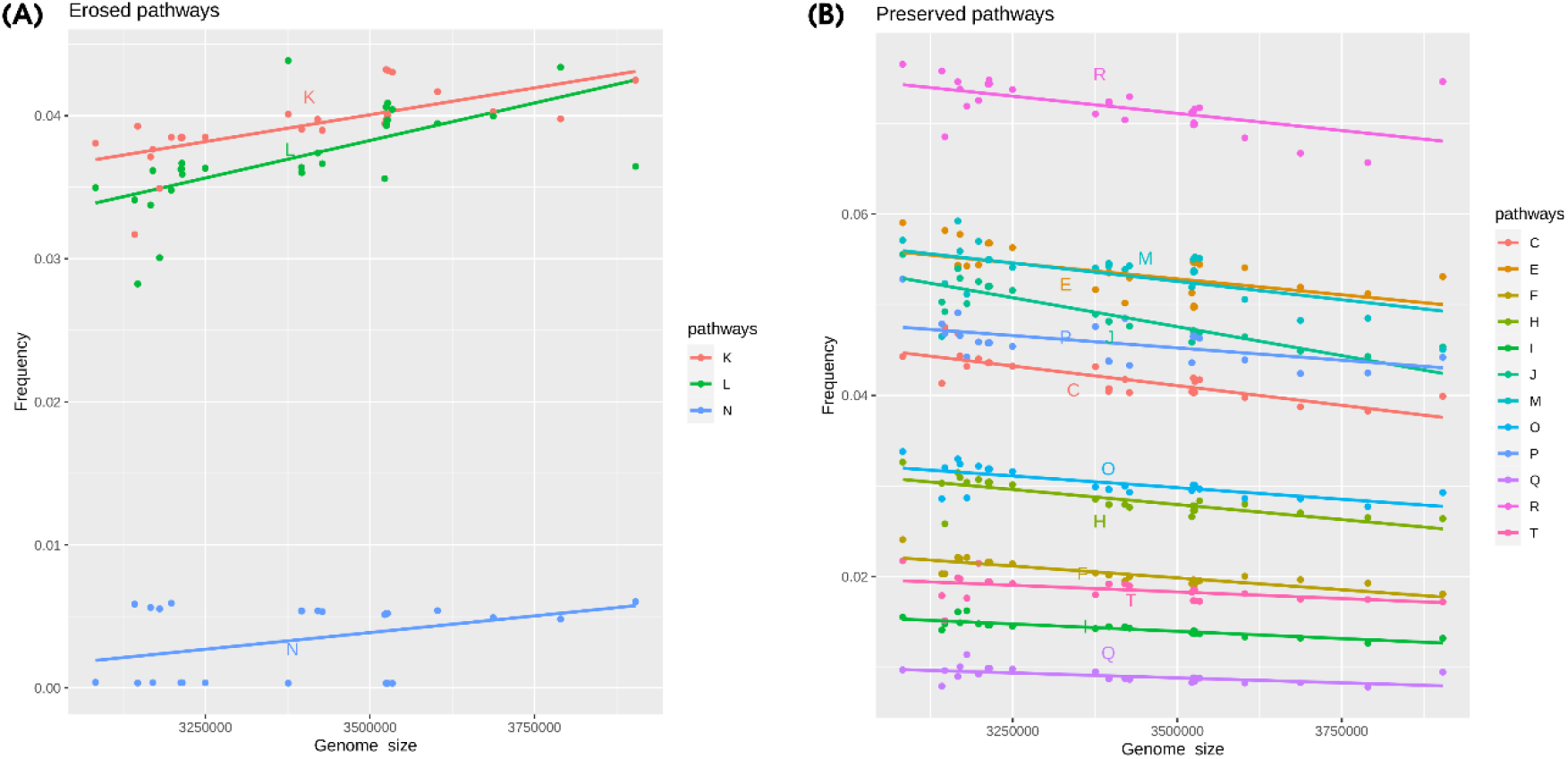
Pathways erosion. For each of the 30 *Asaia* spp. strains included in the study, the proportion of genes belonging to each Cluster of Orthologous Group (COG) were computed as the ratio between the number of genes belonging to a specific pathway and the total number of genes. The scatter plots of COG pathways for which was found a significant linear correlation (Spearman test, p-value < 0.05) between pathway proportion and genome size are reported. In the plot (a), the COG pathways showing a positive correlation are reported, in the plot (b) those showing a negative correlation. A positive correlation suggests that a pathway have been subjected to erosion during genome reduction, conversely a negative correlation suggests its preservation.

Interestingly, the erosion patterns of L, K and N pathways are being conserved during the independent genome reduction processes occurring across the different *Asaia* strains (Fig. 4). These results show that genome erosion pathways follow a conserved stream, likely driven by metabolic constraints. It is also interesting to note that all the *Asaia* strains harbour the *recA* gene, usually lost in highly reduced genomes (20, 21). Furthermore, three plants-derived strains show a slightly different pattern of erosion regarding the two pathways with the highest percentage of eroded genes (Fig. 3). This suggests that, despite the two pathways being associated with cellular/genomic stability, the environment still plays an important role in the definition of the genomic erosion process. This different erosion pattern could be a consequence of a strategy to evade Muller’s ratchet phenomenon (22). As proposed by Naito & Pawlowska (23), the preservation of genes involved in genome stability (e.g. recombination genes) can reduce the accumulation of disadvantageous mutations due to the bottlenecks associated with mother to offspring bacterial transmission (22). We can speculate that *Asaia* strains associated with insect experience bottlenecks more frequently than those living in the environment.

Moreover, the heatmap that takes into account only the presence of pseudogenes, clearly demonstrates that some genes are eroded in many organisms (Fig. 4 and Fig. 5). It also shows that each *Asaia* strain isolated from plants, phylogenetically distant from those isolated from insects (Fig. 1), belongs to a different cluster denoting differentiated reduction patterns. These results suggest a parallel and convergent reduction process of *Asaia* strains from insects.

**Fig. 4.**
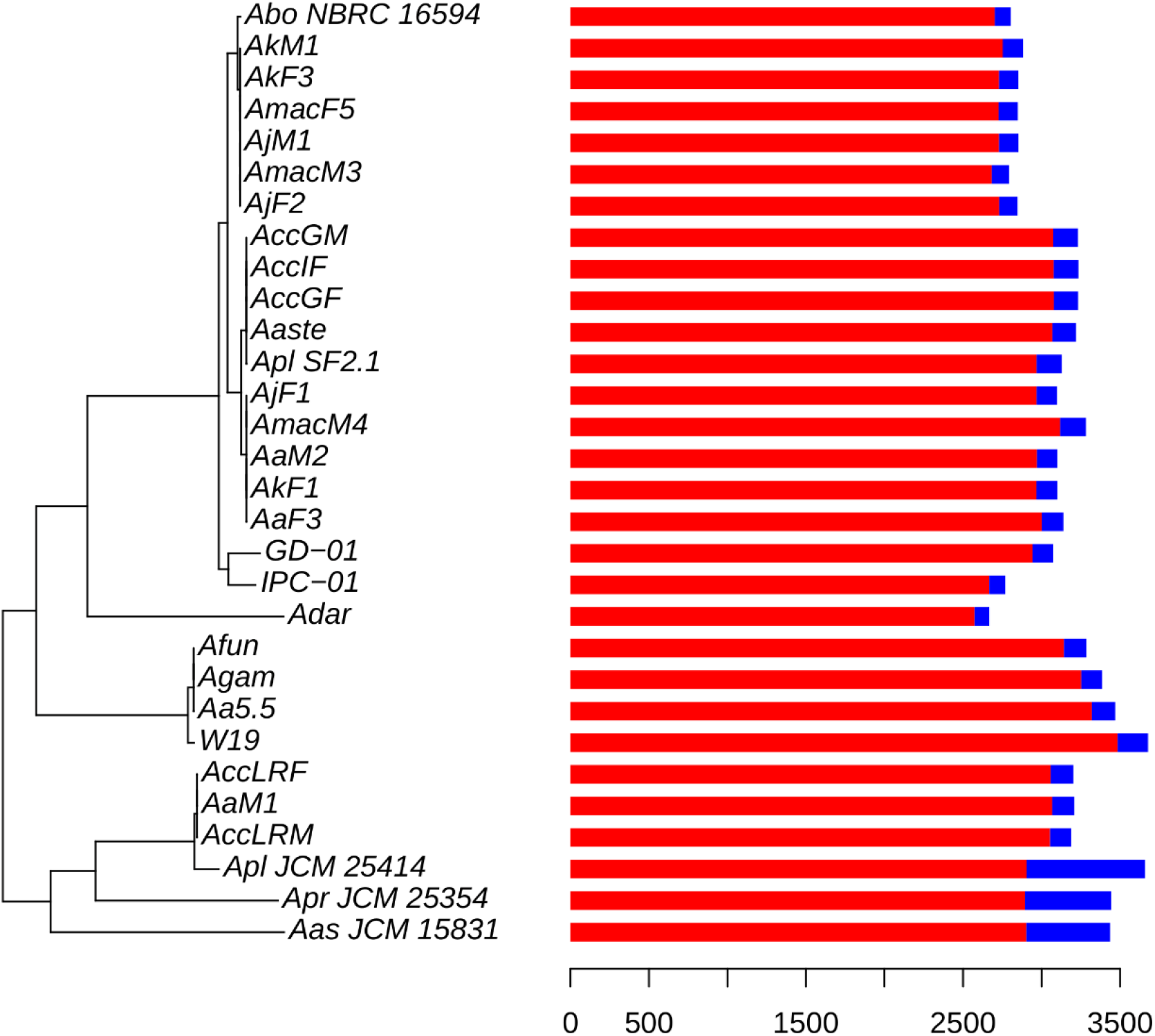
Eroded and entire genes among *Asaia* spp. genomes. Maximum Likelihood phylogenetic tree obtained using the 612 core genes among the 30 *Asaia* spp. strains included in the study and *Gluconobacter morbifer* strain G707 (outgroup). The tree has been rooted on the outgroup strain that was then removed. The bar plot reports the total number of pseudogenes (in blue) and non-peudogenes genes (in red).

**Fig. 5.**
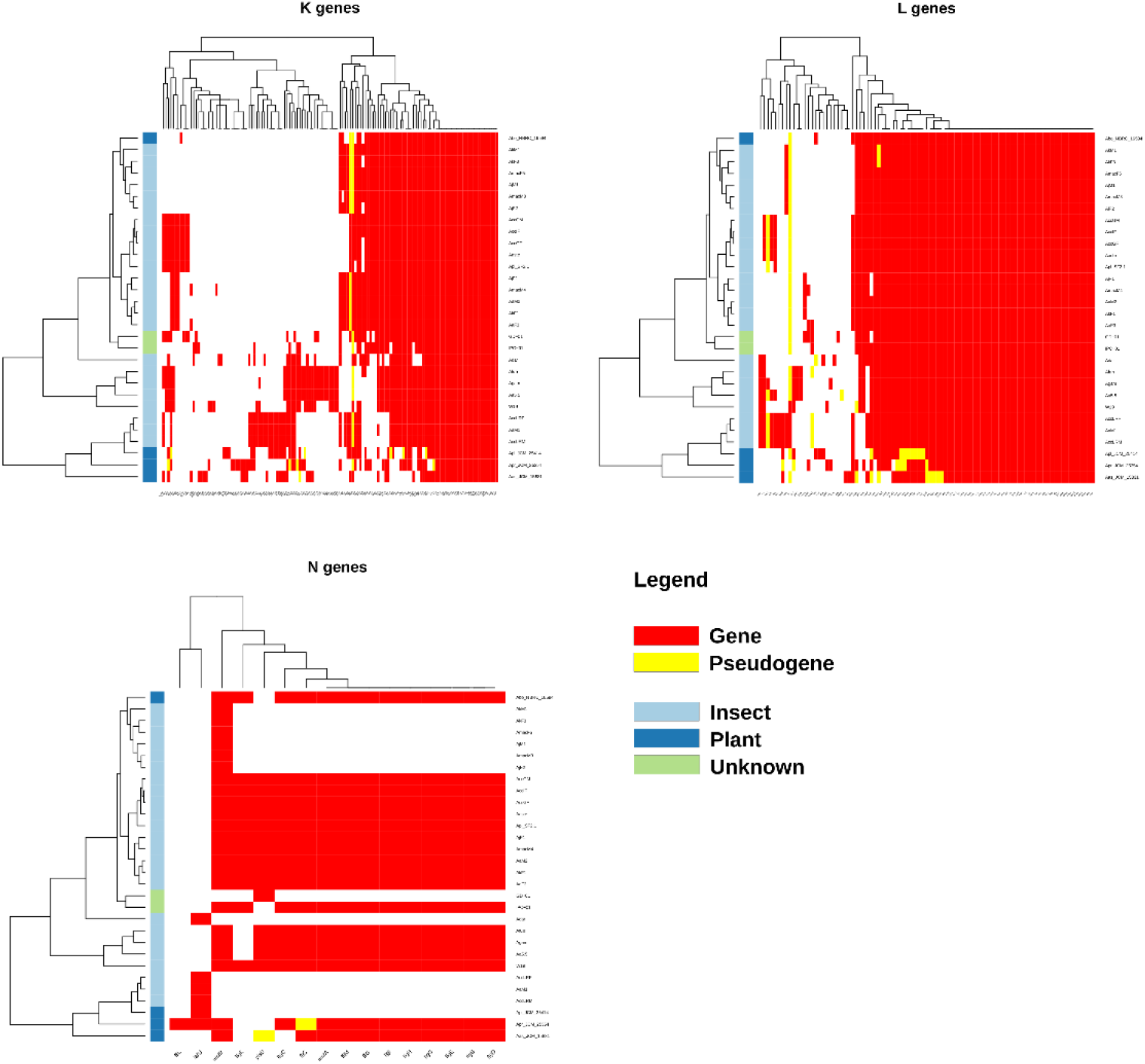
Eroded COG pathways. The heatmap of gene presence/ absence of the eroded COG pathways K (Transcription), L (Replication, recombination and repair) and N (Cell motility) are reported. In each heatmap plot: on the left, the Maximum Likelihood phylogenetic tree of the 30 *Asaia* spp. strains included in the study, in the middle-coloured bar relative to the strain isolation sources (insect in azure, plant in blue and unknown in green), on the right gene presence absence information is reported (present genes in red, pseudogenes in orange and absent in white). In each heatmap, the genes are clustered as obtained using hclust R function.

We have produced an additional heatmap where the frequencies of pseudogenes belonging to each COGs cluster are reported for each organism (Fig. 6). This analysis reveals a substantial convergence in the reduction, excepting for the three *Asaia* isolates from plants that tend to reduce some pathways more than others. Interestingly, the frequency of pseudogenes of the Transcription pathway (K) is not very high, as well as occurs in the Cell Motility pathway (N). The Replication, recombination and repair (L pathway, on the other hand, is one of the richest in pseudogenes. This could be due to the fact that the erosion of the K and N pathways is recent and therefore we observe only a limited number of pseudogenes. On the other hand, the erosion of the Replication, recombination and repair (L) pathway would seem a more gradual process and therefore to be considered still ongoing. All this seems in accordance with the hypothesis of escape from the Muller’s ratchet: in order not to accumulate deleterious mutations, the pseudogenes preserve the path of recombination while eroding the other two pathways much faster. Interestingly, while the erosion of the L pathway (recombination) is fairly conserved among the different lineages (Fig. 6), this does not apply to the K pathway, which shows blocks of different genes conserved through the different lineages. This means that the reduction of the transcriptional apparatus can reach different stable “configurations”.

**Fig. 6.**
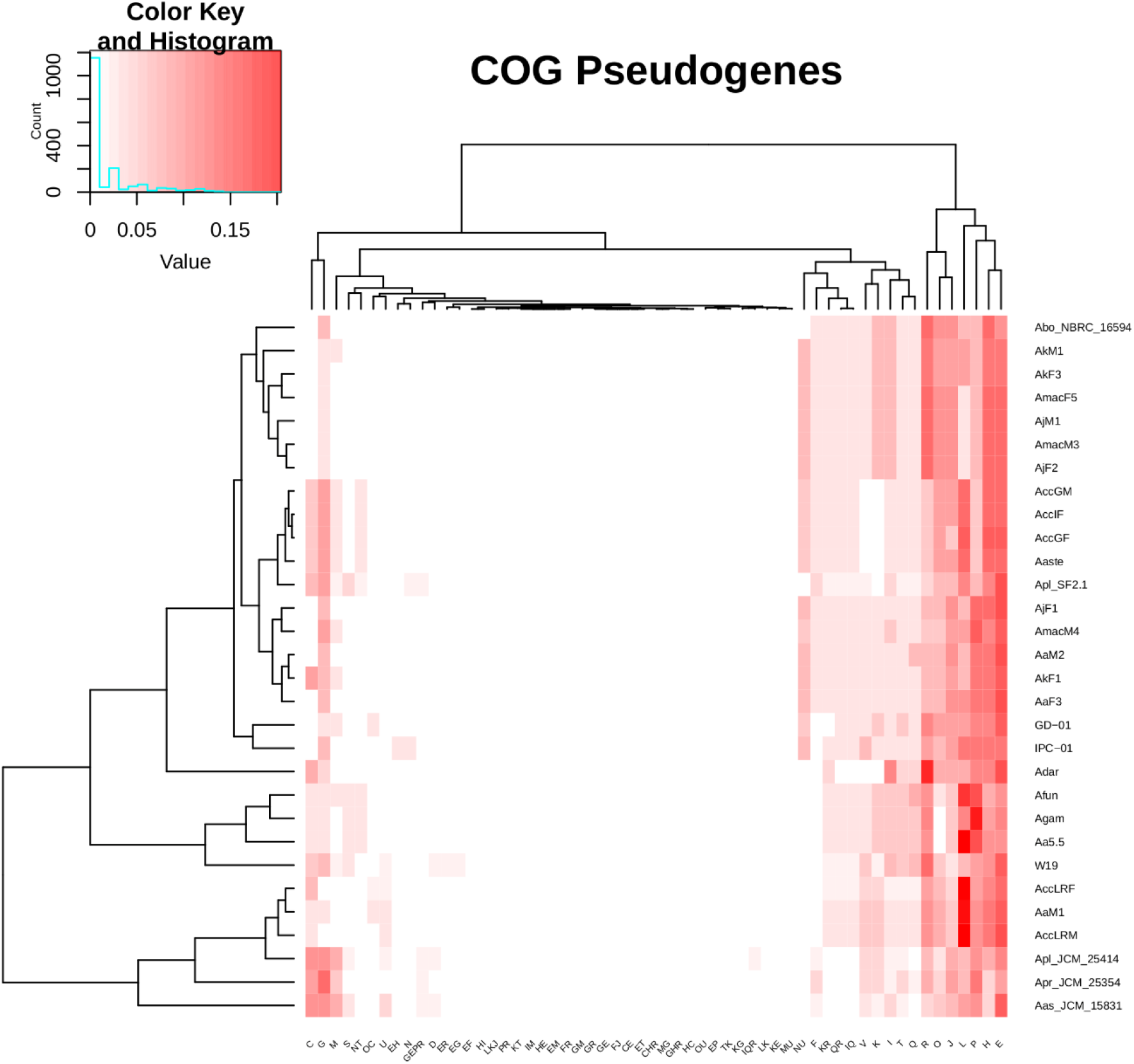
Proportion of pseudogenes among COG pathways. On the left the Maximum Likelihood phylogenetic tree obtained using core genes was rooted on the outgroup (then removed). On the right, colours represent the proportion of pseudogenes present in each COG pathway in each strain. For each strain, the proportion is computed as the ratio between the number of pseudogenes present in a COG pathway and the total number of pseudogenes.

Thus, we focused on some genes and/or metabolic patterns we had focused on in our previous studies because maintained across different *Asaia* strains (5,13).

In the first instance we turned our attention to the pyrethroid hydrolase (PH) gene that regulates pyrethroid degradation, whose mosquito counterpart is known to play a role in pyrethroid resistance. We therefore hypothesized what role it could play in the biology of *Asaia*. The gene coding for PH was found to be present in all but one *Asaia* strains, not only in those isolated from insects. Interestingly this gene is not present in *Asaia* isolated from *An. darlingi* that is the smallest bacterial genome among all those analyzed in this study. The consistency between the phylogenetic analysis of the gene and the phylogenomic analysis reveals that this gene is probably an ancestral character of the genus *Asaia* (Fig. 7). Accordingly, it is possible to hypothesize different roles of PH in the biology of the bacterium. In fact, given the recognized role of PH in the protection against pyrethroids (which are also present in nature as they are produced by some plants), the retention of the PH gene could be due to its own protection or to indirectly protect the insect host from pyrethroids toxicity.

**Fig. 7.**
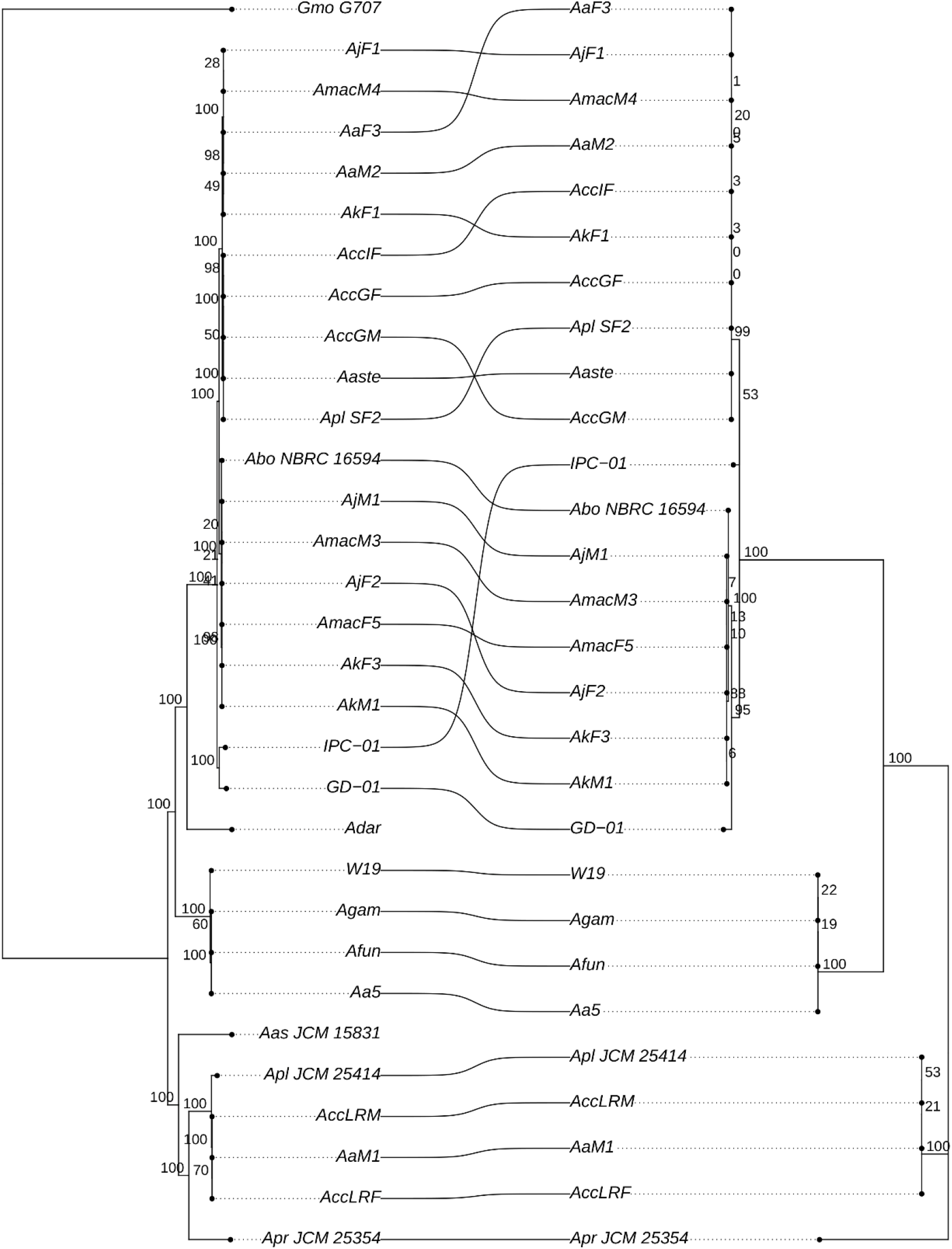
Comparison among *Asaia* spp. species tree and pyrethroid hydrolase phylogenetic tree. On the left the Maximum Likelihood phylogenetic tree obtained using 612 core genes shared among the 30 *Asaia* spp. strains and the outgroup (*Gluconobacter morbifer* strain G707). On the right, the Maximum Likelihood phylogenetic tree reconstructed on the basis of the pyrethroid hydrolase aminoacidic sequence. Bootstrap supports are reported for both trees. Strains shared among the two trees are connected by curved lines. The absence of the lines relative to Adar strain and the outgroup (Gmo G707) indicates the absence of the pyrethroid hydrolase gene in their genomes.

Consequently, the *Asaia* strain isolated from *An. darlingi* missing the PH gene, stands out as an ideal candidate to test this hypothesis, also and above all in light of a recent study aimed to evaluate the susceptibility of *An. darlingi* and *An. marajoara* to pyrethroids used by the National Malaria Control Program in Brazil. This study showed that no resistance was recorded for *An. darlingi* to cypermethrin, deltamethrin, and alpha-cypermethrin, while diagnostic doses estimated for *An. marajoara* indicate that this species requires attention (24). It is important to remark that changes in Anopheles vector competence for malaria parasites have been linked to insecticide resistance status. In fact, insecticide exposure is detrimental for *Plasmodium* in the midgut lumen and is more than likely that additional resistance mechanisms impact parasite development within the vector host (25). It is worth to mention that there are already several examples of insect symbionts that contribute in whole or in part to the lower resistance of the host to the insecticide. In 2012 Kikuchi and collaborators demonstrated that the fenitrothion-degrading *Burkholderia* strains establish a specific and beneficial symbiosis with the stinkbugs and confer a resistance of the host insects against fenitrothion (26). More recently, it has proven that *Hamiltonella defensa* can impact on host aphid susceptibility to some insecticides. In fact, compared with *Hamiltonella-free* aphid clones, *Hamiltonella-* infected aphid clones exhibit lower sensitivity to most of the tested insecticides at low concentrations (27). More generally, although the mechanisms of resistance to insecticides are diversified, detoxification from insecticides mediated by symbionts within insect pests and vectors is a trend that has now become clear, and which is configured as an emerging problem in control of these insects (28).

The second group of genes that has been considered is the one responsible for motility. Our previous study has shown that the genome of the *Asaia* strain isolated from *An. darlingi* had lost a large group of genes involved in the flagellar machinery (i.e., flagella biosynthesis proteins, flagella basal-body rod proteins, flagella motor rotation proteins) (5). In particular, we identified both mobile and not-mobile strains of *Asaia* (see Table S1 in the supplemental material). Moreover, it is worth mentioning that “swimming motility” has been implicated in gut colonization of several hosts, influencing also pathogen infection in mosquitoes (29).

Mobile strains of *Asaia* have larger genomes compared to the immobile strains (Wilcoxon test, p-value = 0.001). Nonetheless, the analysis showed that genes involved in twitching, pili and biofilm production are present in both mobile and immobile strains. These results are greatly supporting of the notion according to which the acquisition of environmental symbionts often necessitates active migration and colonization by the symbionts through motility and chemotaxis (30).

We had already proposed the insect symbiont *Asaia* as a novel model to study genome reduction dynamics, within a single bacterial taxon, evolving in a common biological niche. In contrast to what usually happens in genomic comparison studies for the study of genomic reduction involving genomes substantially different from each other in size (31), in the present study, differences in genomic size fluctuate in a 0.5Mb range. Conserved patterns between independent lines in this dataset suggest that genomic reduction may be an ongoing biological process within the *Asaia* genus, thus confirming this insect symbiont as a model for genome reduction studies.

This study pinpoints the importance of the environment in the definition of the genomic erosion process, selecting and preserving those genes involved in genome stability. On the other hand, our study highlights the role of specific genes in the host biology. In particular, the identification of the pyrethroid hydrolase gene present in almost all the *Asaia* strains isolated beside the one from the South American malaria vector *An. darlingi*, for which resistance to pyrethroids has never been reported, opens the way to a possible involvement of bacteria symbionts in determining the resistance to insecticides developed by insect vectors and insect pests. If confirmed, this involvement would require a revision of the control methods of insect vectors and pest, currently in use.

## Materials and Methods

### *Asaia* strains collection

*Asaia* strains were obtained from different mosquito species and from three different *C. capitata* populations. *Asaia* strains details are listed and described in *SI Appendix*, Table S1. Bacteria isolations was performed as reported in Favia et al (7). Briefly, the insect surface bodies were sterilized washing twice in PBS and Ethanol 70%. After homogenization, *Asaia* was isolated from a pool of five insects by a pre-enrichment step in ABEM liquid medium (2% sorbitol; 0.5% peptone; 0.3% yeast extract; 100 μg/ml actidione, pH 3.5) and hence plated in agarized medium (2% glucose; 0.5% ethanol; 0.8% yeast extract; 0.7% carbonate calcium; 1.2% agar; pH 7). *Asaia* colonies were identified by their morphology and ability to produce carbonate dissolution haloes in agar plates. Identification was then confirmed by PCR using 1492Rev-27For oligonucleotides targeting 16S rRNA gene and sequencing by sanger methods (Eurofins, Germany). *Asaia* motility test was conducted using the hanging drop procedure. Each bacterial strain was grown on GLY agar (yeast extract 1%, glycerol 2.5%, agar 2%, pH 5) for 48 h at 30 C. After picking a single colony from each plate, bacteria were re-suspended in a drop of 1X PBS and then observed under 100 optical microscopy. Cell motility was assessed by observing multiple areas of the same slide within three biological replicates (see Table S1 in the supplemental material). *Asaia* genome sequence of environmental strains were recovered by PATRIC database (see Table S3 in the supplemental material).

### DNA extraction and genome sequencing

The 17 *Asaia* spp. strains were subjected to whole DNA extraction and Whole Genome Sequencing (WGS). Genomic DNA was extracted using the QIAamp DNA minikit (Qiagen) following the manufacturer’s instructions and Whole Genome Sequencing was performed using Illumina Miseq platform with a 2 x 250 paired-end run after Nextera XT library preparation (Illumina Inc., San Diego, CA).

### Genome retrieving

All the genome assemblies and genomic reads of *Asaia* spp. strains available on public databases on 1^st^ March 2020 were retrieved, for a total of eight strains genome assemblies (retrieved from NCBI and PATRIC) and five strains short-reads data files (retrieved from Sequence Read Archive, SRA). Furthermore, the genome assembly of *Gluconobacter morbifer* str. Gmo_G707, included as outgroup in the phylogenomic analyses, was retrieved from NCBI. For more details, see Table S3 in the supplemental material.

### Genome assembly

The paired-end reads obtained from the 17 *Asaia* strains and the five reads sequences retrieved from SRA were subjected to quality check using FastQC tool (32) and then assembled using SPAdes 3.13 (33).

### Average Nucleotide Identity (ANI) based genomes clustering

The pairwise Average Nucleotide Identity values among the 31 genomes included in the study were computed using OrthoANI tool (34). Genomes were then clustered using a cut-off threshold of 95% (35).

### Genome annotation and orthologous analysis

All the 31 genome assemblies included in the study were subjected to Open Reading Frame (ORF) calling and annotation using PROKKA (36). Amino acid sequences obtained from PROKKA for all the genome assemblies were then used for orthologous analysis using OrthoMCL (37). The output of the orthologous analysis was then refined using the annotation information: two sequences were included in the same orthologous group if they belonged to the same OrthoMCL orthologous group and they have the same PROKKA annotation.

### Identification of pseudogenes

For each refined orthologous group, the sequences were retrieved, aligned and analyzed to identify pseudogenes. We classified as pseudogenes the sequences containing one (or more) internal stop codon, one (or more) frame shift or those for which > 30% of the bases in the alignment are gaps. More in detail, for each group, the amino acid sequences were aligned using MUSCLE (38) and the obtained alignment was used to obtain the relative codon-based nucleotide alignment. The obtained nucleotide alignments were then analysed using MACSE 2.03 (39) to identify internal stop codons and frame shifts. The percentage of gap positions for each sequence of each nucleotide alignment was obtained using a Perl script.

### Phylogenomic analysis

Core genes were identified as pseudogenes-free orthologous groups containing one gene sequence from each of the 31 genome assemblies included in the study. Amino acid alignments of core genes were retrieved, concatenated and subjected to Maximum Likelihood (ML) phylogenetic analysis. The ML phylogenetic analysis was conducted using RAxML 8 (40) using the LG+I+G+F model, selected using ProtTest3 (41).

### Principal Coordinates Analysis (PCoA)

The output of orthologous analysis was converted in a gene presence/absence matrix and PCoA analysis was performed out using R.

### Pathways erosion analysis

To investigate relationship between pathway erosion and genome size we annotated all the amino acid sequences of the 30 *Asaia* strains genomes by BLASTP search against the Clusters of Orthologous Groups of proteins (COGs) database. Then, for each genome, we computed the proportion of genes of each COG pathway as the ratio between the number of the genes belonging to each COG pathway and the total number of genes. Then, for each COG pathway, we computed the linear regression analysis between the pathway proportion and genome size considering the 30 *Asaia* strains included in the study.

### Pyrethroid hydrolase vs species trees co-cladogenesis analysis

The amino acid alignment of the pyrethroid hydrolase gene was retrieved from output of the previous analyses (see above). The alignment was subjected to ML phylogenetic using RAxML 8 (40) with the LG+G+F model, selected using ProtTest3 (41). The topologies of the pyrethroid hydrolase tree and the species tree obtained by phylogenomic analysis (see above) were then graphically compared using R.

## Data availability

All the reads related to the new genome sequencing described in this study has been deposited in the European Nucleotide Archive (ENA) and accession numbers are pending.

## Acknowledgments

This work was supported by PRIN2015JXC3JF from Italian Ministry for University Research (MIUR), FAR2019 from University of Camerino to GF. FC and CB work was supported by MIUR-PRIN grant n. 2017J8JR57. We thank Professor Valerio Napolioni for his valuable comments on the manuscript.

## Author Contributions

All authors read and approved the manuscript. G.F. conceived the work. F.C., C.D., A.C., G.G., F.G., G.C., F.M., P. Ro., carried out sample preparation, sequencing, genome assembly and/or gene finding and annotation. F.C., and G.F., analyzed Phyretroid Hydrolase sequence data. F.C., C.D., A.C., A.P., I.R., M.V.M., P. Ri., C.B. and G.F. performed data analyses. G.F., F.C., C.D., and C.B. analyzed data and wrote the manuscript.

## Supplemental Material

Table S1: *Asaia* strains information

Table S2: Pathways linear regression analysis information

Table S3: Genome information and statistics

Fig. S1: **Maximum Likelihood phylogenetic tree including multiple outgroup species**. Maximum likelihood phylogenetic tree including the study strains of the genus *Asaia* and 12 strains of four genera belonging to the Acetobacteraceae family. The phylogenetic analysis has been perfomed on a concatenate of 356 core genes. These core genes were selected as follows: present in single copy in all the genomes and without frame shifts or stop codons in the alignment. Bootstrap supporting values are reported on the tree branches.

## References

1. Margulis L, Fester R. 1991. Symbiosis as a Source of Evolutionary Innovation: Speciation and Morphogenesis. The MIT Press, Cambridge, MA eds.

2. Moran NA, Bennett GM. 2014. The Tiniest Tiny Genomes. Annu Rev Microbiol 68: 195–215.

3. McCutcheon JP, Moran NA. 2012. Extreme genome reduction in symbiotic bacteria. Nat Rev Microbiol 10: 13–26.

4. Nicks T, Rahn-Lee L. 2017. Inside Out: Archaeal Ectosymbionts Suggest a Second Model of Reduced-Genome Evolution. Front Microbiol 8:384.

5. Alonso DP, Mancini MV, Damiani C, Cappelli A, Ricci I, Alvarez MVN, Bandi C, Ribolla PEM, Favia G. 2019. Genome Reduction in the Mosquito Symbiont *Asaia*. Genome Biol Evol. 11: 1–10.

6. Crotti E, Damiani C, Pajoro M, Gonella E, Rizzi A, Ricci I, Negri I, Scuppa P, Rossi P, Ballarini P, Raddadi N, Marzorati M, Sacchi L, Clementi E, Genchi M, Mandrioli M, Bandi C, Favia G, Alma A, Daffonchio D. 2009. *Asaia*, a versatile acetic acid bacterial symbiont, capable of cross-colonizing insects of phylogenetically distant genera and orders. Environ Microbiol 11: 3252–64

7. Favia G, Ricci I, Damiani C, Raddadi N, Crotti E, Marzorati M, Rizzi A, Urso R, Brusetti L, Borin S, Mora D, Scuppa P, Pasqualini L, Clementi E, Genchi M, Corona S, Negri I, Grandi G, Alma A, Kramer L, Esposito F, Bandi C, Sacchi L, Daffonchio D. 2007. Bacteria of the genus *Asaia* stably associate with *Anopheles stephensi*, an Asian malarial mosquito vector. Proc Natl Acad Sci U S A. 104: 9047–9051.

8. Damiani C, Ricci I, Crotti E, Rossi P, Rizzi A, Scuppa P, Capone A, Ulissi U, Epis S, Genchi M, Sagnon N, Faye I, Kang A, Chouaia B, Whitehorn C, Moussa GW, Mandrioli M, Esposito F, Sacchi L, Bandi C, Daffonchio D, Favia G. 2010. Mosquito-bacteria symbiosis: the case of *Anopheles gambiae* and *Asaia*. Microb Ecol 60: 644–654.

9. Chen S, Zhang D, Augustinos A, Doudoumis V, Bel Mokhtar N, Maiga H, Tsiamis G, Bourtzis K. 2020. Multiple Factors Determine the Structure of Bacterial Communities Associated with *Aedes albopictus* Under Artificial Rearing Conditions. Front Microbiol. 11: 605.

10. De Freece C, Damiani C, Valzano M, D’Amelio S, Cappelli A, Ricci I, Favia G. 2014. Detection and isolation of the α-proteobacterium *Asaia* in *Culex mosquitoes*. Med Vet Entomol 28: 438–442

11. Bassene H, Niang EHA, Fenollar F, Doucoure S, Faye O, Raoult D, Sokhna C, Mediannikov O. 2020. Role of plants in the transmission of *Asaia* sp., which potentially inhibit the *Plasmodium* sporogenic cycle in *Anopheles* mosquitoes. Sci Rep 10: 7144.

12. Chouaia B, Rossi P, Epis S, Mosca M, Ricci I, Damiani C, Ulissi U, Crotti E, Daffonchio D, Bandi C, Favia G. 2012. Delayed larval development in *Anopheles mosquitoes* deprived of *Asaia* bacterial symbionts. BMC Microbiol 12: S2.

13. Mancini MV, Damiani C, Short SM, Cappelli A, Ulissi U, Capone A, Serrao A, Rossi P, Amici A, Kalogris C, Dimopoulos G, Ricci I, Favia G. 2020. Inhibition of *Asaia* in Adult Mosquitoes Causes Male-Specific Mortality and Diverse Transcriptome Changes. Pathogens 9: 380.

14. Cappelli A, Damiani C, Mancini MV, Valzano M, Rossi P, Serrao A, Ricci I, Favia G. 2019. Asaia Activates Immune Genes in Mosquito Eliciting an Anti-*Plasmodium* Response: Implications in Malaria Control. Front Genet 10: 836.

15. Epis S, Varotto-Boccazzi I, Crotti E, Damiani C, Giovati L, Mandrioli M, Biggiogera M, Gabrieli P, Genchi M, Polonelli L, Daffonchio D, Favia G, Bandi C. 2020. Chimeric symbionts expressing a *Wolbachia* protein stimulate mosquito immunity and inhibit filarial parasite development. Commun Biol 3: 105.

16. Li F, Hua H, Ali A, Hou M. 2019. Characterization of a Bacterial Symbiont *Asaia* sp. in the White-Backed Planthopper, *Sogatella furcifera*, and Its Effects on Host Fitness. Front Microbiol 10: 2179.

17. White IM, Elson-Harris M. 1992. Fruit flies of economic significance: their identification and bionomics. International Institute of Entomology, London, United Kingdom.

18. Bhatt P, Huang Y, Zhan H, Chen S. 2019. Insight Into Microbial Applications for the Biodegradation of Pyrethroid Insecticides. Front Microbiol 10: 1778.

19. Jain C, Rodriguez-R LM, Phillippy AM, Konstantinidis KT, Aluru S. 2018. High throughput ANI analysis of 90K prokaryotic genomes reveals clear species boundaries. Nat Commun. 9(1): 5114.

20. Klasson L, Andersson SGE. 2006. Strong asymmetric mutation bias in endosymbiont genomes coincide with loss of genes for replication restart pathways. Mol Biol Evol 23: 1031–1039.

21. Comandatore F, Cordaux R, Bandi C, Blaxter M, Darby A, Makepeace BL, Montagna M, Sassera D. 2005. Supergroup C *Wolbachia*, mutualist symbionts of filarial nematodes, have a distinct genome structure. Open Biol 5: 150099.

22. Moran NA. 1996. Accelerated evolution and Muller’s rachet in endosymbiotic bacteria. Proc Natl Acad Sci USA 93: 2873–2878.

23. Naito M, Pawlowska TE. 2016. Defying Muller’s Ratchet: Ancient Heritable Endobacteria Escape Extinction through Retention of Recombination and Genome Plasticity. mBio 7: e02057–15.

24. Galardo AKR, Póvoa MM, Sucupira IMC, Galardo CD, La Corte Dos Santos R. 2015. *Anopheles darlingi* and *Anopheles marajoara* (Diptera: Culicidae) Susceptibility to Pyrethroids in an Endemic Area of the Brazilian Amazon. Rev Soc Bras Med Trop 48: 765–769.

25. Minetti C, Ingham VA, Ranson H. 2020. Effects of insecticide resistance and exposure on *Plasmodium* development in *Anopheles* mosquitoes. Curr Opin Insect Sci 39: 42–49.

26. Kikuchi Y, Hayatsu M, Hosokawa T, Nagayama A, Tago K, Fukatsu T. 2012. Symbiont-mediated insecticide resistance. Proc Natl Acad Sci U S A. 109(22): 8618–8622.

27. Li X, Liu Q, Liu H, Bi H, Wang Y, Chen X, Wu N, Xu J, Zhang Z, Huang Y, Chen H. 2020. Mutation of doublesex in *Hyphantria cunea* results in sex-specific sterility. Pest Manag Sci. 76(5):1673–1682.

28. Blanton AG, Peterson BF. 2020. Symbiont-Mediated Insecticide Detoxification as an Emerging Problem in Insect Pests. Front Microbiol 11:547108.

29. Bando H, Okado K, Guelbeogo WM, Badolo A, Aonuma H, Nelson B, Fukumoto S, Xuan X, Sagnon N, Kanuka H. 2013. Intra-specific diversity of *Serratia marcescens* in *Anopheles* mosquito midgut defines *Plasmodium* transmission capacity. Sci Rep 3:1641.

30. Raina JB, Fernandez V, Lambert B, Stocker R, Seymour JR. 2019. The Role of Microbial Motility and Chemotaxis in Symbiosis. Nat Rev Microbiol 17: 284–294.

31. Manzano-Marín A, Latorre A. 2016. Snapshots of a Shrinking Partner: Genome Reduction in *Serratia symbiotica*. Sci Rep 6:32590.

32. Andrews S. 2010. FastQC: A Quality Control Tool for High Throughput Sequence Data. Available online at:http://www.bioinformatics.babraham.ac.uk/projects/fastqc/.

33. Bankevich A, Nurk S, Antipov D, Gurevich AA, Dvorkin M, Kulikov AS, Lesin VM, Nikolenko SI, Pham S, Prjibelski AD, Pyshkin AV, Sirotkin AV, Vyahhi N, Tesler G, Alekseyev MA, Pevzner PA. 2012. SPAdes: a new genome assembly algorithm and its applications to single-cell sequencing. J Comput Biol 19: 455–477.

34. Lee I, Kim YO, Park SC, Chun J. 2016. OrthoANI: an improved algorithm and software for calculating average nucleotide identity. Int J Syst Evol Micr 66: 1100–1103.

35. Jain C, Rodriguez-R LM, Phillippy AM, Konstantinidis KT, Aluru S. 2018. High throughput ANI analysis of 90K prokaryotic genomes reveals clear species boundaries. Nat Comm 9: 1–8.

36. Seemann T. 2014. Prokka: rapid prokaryotic genome annotation. Bioinformatics 30: 2068–2069.

37. Li L, Stoeckert CJ, Roos DS. 2003. OrthoMCL: identification of ortholog groups for eukaryotic genomes. Genome Res 13: 2178–2189.

38. Edgar RC. 2004. MUSCLE: multiple sequence alignment with high accuracy and high throughput. Nucleic Acids Res. 32: 1792–1797.

39. Ranwez V, Harispe S, Delsuc F, Douzery EJ. 2011. MACSE: Multiple Alignment of Coding Sequences accounting for frameshifts and stop codons. PloS One 6: e22594.

40. Stamatakis A. 2014. RAxML Version 8: A tool for Phylogenetic Analysis and Post-Analysis of Large Phylogenies. Bioinformatics 30: 1312–1313.

41. Darriba D, Taboada GL, Doallo R, Posada D. 2011. ProtTest 3: fast selection of best-fit models of protein evolution. Bioinformatics 27: 1164–1165.

